# Ribosome stalling facilitates chloroplast targeting of nuclear-encoded proteins

**DOI:** 10.64898/2026.03.25.714110

**Authors:** Ryosuke Kowada, Shintaro Iwasaki, Yukihide Tomari, Hiro-oki Iwakawa

**Affiliations:** Department of Computational Biology and Medical Sciences, Graduate School of Frontier Sciences, The University of Tokyo, Bunkyo-ku, Tokyo, Japan; Laboratory of RNA Function, Institute for Quantitative Biosciences, The University of Tokyo, Bunkyo-ku, Tokyo, Japan; Department of Life Science, College of Science, Rikkyo University, Toshima-ku, Tokyo, Japan; RNA Systems Biochemistry Laboratory, Pioneering Research Institute, RIKEN, Wako, Saitama, Japan

**Author notes:** Corresponding author (H.-o.I.) and (Y.T.).

## Abstract

Ribosomes translate mRNAs with variable elongation rates and frequently undergo transient stalling. Although ribosome stalling is known to regulate protein quality control and translational dynamics, its transcriptome-wide landscape and biological significance in plants remain largely unexplored. Here, using disome profiling in *Arabidopsis thaliana*, we generated a high-resolution map of ribosome stalling sites across the transcriptome and uncovered a marked enrichment on mRNAs encoding chloroplast-targeted proteins, particularly those involved in photosynthesis. We further found that ribosome stalling preferentially occurs when transit peptides emerge from the ribosome exit tunnel. Functional assays demonstrated that deletion of the stalling region significantly reduces chloroplast targeting efficiency. Together, our findings identify ribosome stalling as a regulatory layer that promotes efficient targeting of nuclear-encoded proteins to chloroplasts.

## Introductory paragraph

Ribosomes translate mRNAs at non-uniform rates and can transiently pause during elongation^1–3^. Such pauses, termed ribosome stalling, contribute to protein quality control, translational regulation, and other cellular processes^4–6^. Stalling often leads to ribosome collisions, generating disome and trisome complexes that serve as molecular signatures of translational slowdown^7–9^. These events can be triggered by intrinsic features of the nascent peptide^10–15^, including specific sequence motifs and positively charged residues, as well as by extrinsic obstacles on mRNAs^16–18^, such as RNA-induced silencing complex (RISC) composed of microRNAs and Argonaute proteins. While the physiological roles of ribosome stalling have been extensively studied in yeast and mammalian systems^4,19,20^, its biological significance in plants remains largely unexplored, despite the unique intracellular organization and organelle complexity that distinguish plant cells from other eukaryotes.

## Main

Here, we applied disome profiling in plants—a specialized version of the conventional ribosome profiling (monosome profiling)^21,22^ designed to capture the approximately twice-long footprints generated by collided ribosomes^2,9,23–25^. Using this approach, we mapped disome footprints across the transcriptome (Fig. 1a) in three-day-old *Arabidopsis thaliana* seedlings. Mapping disome reads to nuclear-encoded transcripts revealed distinct disome footprint length peaks at 54, 58, and 61 nt for cytoplasmic ribosomes (Fig. 1b, upper panel). These longer footprints contrasted sharply with monosome footprints^17,26^, which peaked at 21 and 28 nt (Fig. 1c, upper panel). In addition to reads within CDS, we observed high disome accumulation at the stop codons (Fig. 1d), likely reflecting slow termination and/or ribosome recycling, as reported in other eukaryotes^25,27,28^. In the metagene analysis, the 5ʹ ends of disome footprints were positioned 45 nt upstream of the stop codon (Fig. 1d), consistent with the combined length of a monosome footprint (∼30 nt) and the A-site offset (∼15 nt). Using this offset, we assigned the A-site codons of the leading ribosome in the disomes. Notably, disomes did not accumulate within the 12-nt region upstream of start codons (Fig. 1e), consistent with previous observations in yeast, zebrafish, and humans^9,23,25^. This upstream space likely reflects occupancy by factors associated with the scanning ribosome, suggesting conservation of ribosome scanning mechanisms across eukaryotes. To validate our disome profiling data, we examined genes previously reported to induce ribosome stalling in plants, including *CGS1*^29,30^ and *bZIP60u*^31^, for which stalling sites have been biochemically characterized. In both genes, prominent disome peaks were observed at the expected positions (Fig. 1f, Extended Data Fig. 1). Similarly, on the *TAS3a* transcript, where ribosome stalling is induced by RISC-mediated obstruction of translating ribosomes^17^, sharp disome peaks were detected at the same positions as the monosome peaks (Fig. 1g). These results demonstrate that disome profiling reliably detects ribosome stalling driven by diverse regulatory mechanisms.

**Fig. 1.**
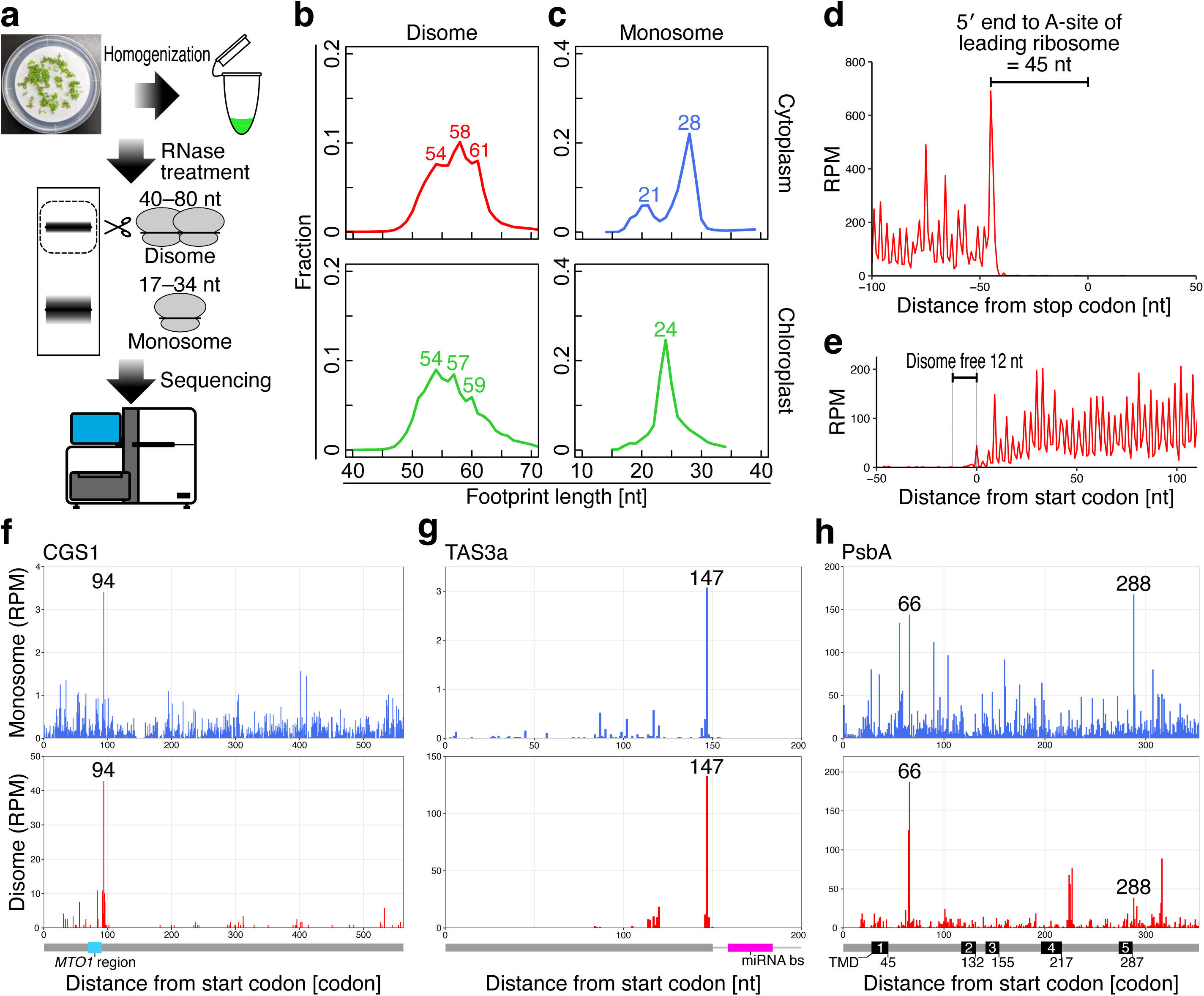
Metagene analysis of disome profiling in plants and benchmarking using established ribosome stalling genes. **a**, Schematic workflow for disome profiling library preparation. **b**, Disome footprint length distribution in 3-day-old *Arabidopsis thaliana* seedlings. Cytoplasmic (upper; red) and chloroplast (lower; green) ribosome footprints are shown. **c**, Monosome footprint length distribution reanalyzed from previous data^17^. Cytoplasmic (upper; blue) and chloroplast (lower; green) ribosome footprints are shown. **d**,**e**, Metagene analysis of 58-nt disome footprints aligned at the 5ʹ ends around the stop codon (**d**) and start codon (**e**). The first nucleotide of the start or stop codon is defined as position 0. **f**,**g**, Footprint distributions along nuclear-encoded transcripts with established ribosome stalling sites: *CGS1* (**f**) and *TAS3a* (**g**). A-site positions are plotted for monosome (upper; blue bars) and the leading ribosome in a disome (lower; red bars). For CGS1, the established stalling site at Ser94 (94) is indicated in both panels. A schematic of the full-length protein is shown below (gray), with MTO1 region highlighted in light blue (**f**). For TAS3a, the ORF is shown (gray line), and the miRNA binding site (miRNA bs) is indicated in magenta (**g**). RPM: reads per million mapped reads. **h**, Footprint distribution along the chloroplast-encoded *PsbA* gene. Panel arrangement follows that of (**f**, **g**). The previously reported stalling site at codon 288 and the prominent peak at codon 66 identified in this study are indicated. A schematic of the full-length protein is shown below (gray), with predicted transmembrane domains (TMDs) represented by black boxes. RPM: reads per million mapped reads.

Disome profiling also captured footprints from chloroplast ribosomes. Mapping reads to chloroplast-encoded transcripts revealed length peaks at 54, 57, and 59 nt (Fig. 1b, lower panel), distinct from chloroplast monosome footprints (Fig. 1c, lower panel; Wakigawa et al.^26^), mirroring the trend observed for cytoplasmic ribosomes. We next examined ribosome stalling in the chloroplast-encoded gene *PsbA*^32,33^, which encodes the D1 reaction center protein of Photosystem II (PSII) and contains five transmembrane domains (TMDs). Previous ribosome profiling^26^ showed abundant monosome accumulation at codon 288, corresponding to the emergence of the fourth TMD from the ribosome exit tunnel. Consistent with this, our monosome profiling data^17^ also displayed a pronounced peak at codon 288 (Fig. 1h). In contrast, disome profiling revealed a prominent stalling site at codon 66, corresponding to the emergence of the first TMD (Fig. 1h). We propose that ribosome stalling at this early TMD may provide time for proper membrane engagement of the nascent peptide. One possible explanation for the absence of disome accumulation at the fourth TMD is that pausing at the first TMD limits the collision of trailing ribosomes at the downstream positions.

Given the robust performance of disome profiling, we next conducted a transcriptome-wide identification of stalling sites on nuclear-encoded transcripts. Stalling sites were defined as nucleotide positions accumulating more than 5% of total disome footprints within a CDS and exceeding 10 RPM. Mapping disome footprints identified 181,069 candidate sites (Fig. 2a), with strong reproducibility between biological replicates even at single-nucleotide resolution (Extended Data Fig. 2a). Of these, 646 sites satisfied the stalling criteria in both replicates, corresponding to 288 genes excluding splice variants. These genes were not biased toward lowly expressed transcripts, as RNA-seq and monosome profiling analyses confirmed substantial RNA abundance and basal translation levels (Extended Data Fig. 2b, c). Together, these results indicate that the detected ribosome stalling events represent reproducible and biologically meaningful translational pauses rather than stochastic sampling noise.

**Fig. 2.**
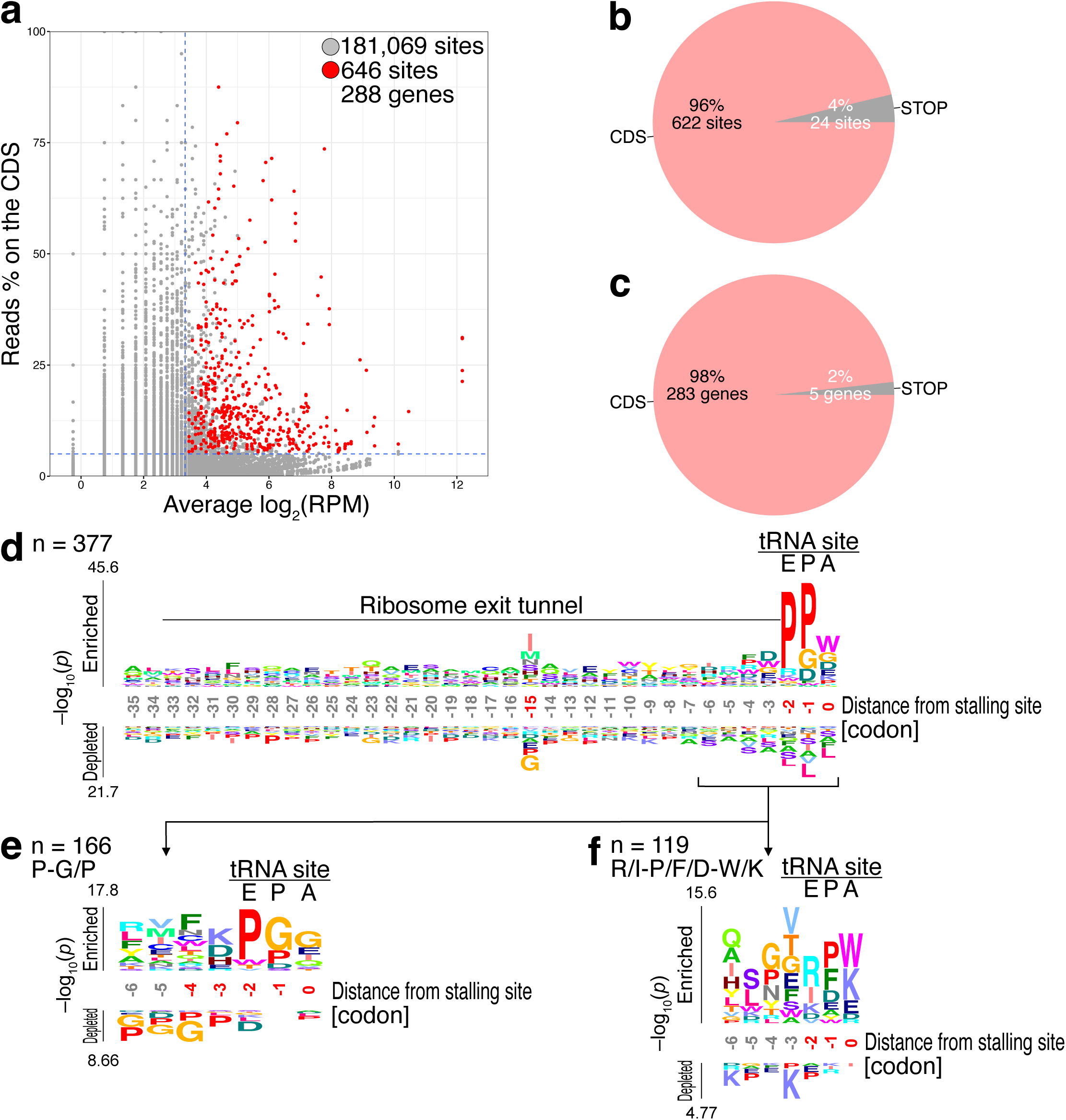
Transcriptome-wide analysis reveals that ribosome stalling tends to occur on mRNAs encoding chloroplast-localized proteins. **a**, Scatter plot of disome footprints mapped to all nuclear-encoded transcripts annotated in TAIR10. Each point represents the first nucleotide of the A-site of the leading ribosome in a disome. Gray dots indicate all identified sites, and red dots indicate sites satisfying the stalling criteria in both replicates. The x-axis represents the mean RPM at the site of the two replicates, and the y-axis represents the ratio of reads at each position to the total reads within each CDS. Horizontal and vertical blue dotted lines indicate the thresholds of 5% and 10 RPM, respectively. **b**,**c**, Pie charts indicating whether sites satisfying the stalling criteria are located within the CDS or at the stop codon. Panel **b** summarizes all candidate sites (622 CDS sites; 24 stop codon sites), and panel **c** summarizes genes containing these stalling sites (283 genes with CDS stalling; 5 genes with stop codon stalling). No gene contains stalling sites in both regions simultaneously. **d**–**f**, Amino acid motif analysis of CDS stalling sites aligned at the A-site of the leading ribosome in a disome. Overall motif analysis (**d**). Sub-cluster analyses based on residues surrounding the tRNA-binding sites (E, P, and A sites) (**e** and **f**). n represents the number of sequences. Sequence logos were generated using *k*pLogo^56^, and significantly enriched positions are highlighted in red (FDR < 0.01).

To define the positional characteristics of the 646 high-confidence stalling sites (288 genes), we first examined their genomic distribution. Over 96% were located within CDSs (622 sites; 283 genes), whereas fewer than 4% occurred at stop codons (24 sites; 5 genes) (Fig. 2b, c), indicating the majority represent previously uncharacterized elongation pauses. We next investigated sequence features associated with CDS-localized sites, restricting the analysis to distinct amino acid sequences to avoid redundancy from splice variants. Motif enrichment analysis revealed a prominent Pro-Pro-Trp motif at the E-P-A sites of the leading ribosome in the disomes (Fig. 2d). To separate the effects of multiple amino acid motifs, we further clustered ribosome stalling sites based on residues surrounding the tRNA-binding site of the leading ribosome. This analysis identified two major stalling motifs: Pro-Gly/Pro at the E-P sites (Fig. 2e) and Arg/Ile-Pro/Phe/Asp-Trp/Lys at the E-P-A sites (Fig. 2f). The former has been shown to induce ribosome stalling via unfavorable interactions with the peptidyl transferase center^10,34,35^, whereas the latter likely impedes elongation through a combination of electrostatic interactions with rRNA^12,23,36^, steric hindrance imposed by bulky residues, and variations in intracellular amino acid usage.

Notably, Gene Ontology (GO) analysis of the 283 genes with stalling sites in CDSs revealed a strong enrichment of photosynthesis-related biological processes, including light harvesting, photochemical reactions, and carbon fixation (Fig. 3a), as well as molecular functions such as NAD^+^/NADP^+^ activity and chlorophyll binding (Fig. 3b). One possibility is that genes with readily detectable disomes simply correspond to highly translated transcripts with abundant monosome footprints. However, GO analysis of the top 10% of transcripts with the highest monosome footprint abundance (4,140 transcripts) showed that their enriched GO terms were largely unrelated to chloroplast functions (Extended Data Fig. 3a, b). Furthermore, when the stalling genes were analyzed against this highly translated background, chloroplast-related GO terms remained prominently enriched (Extended Data Fig. 3c). Thus, the enrichment of chloroplast-related GO terms cannot be explained solely by high translation levels. Consistent with this interpretation, analysis of UniProt annotations revealed that 100 of the 283 genes encode chloroplast-localized proteins (Fig. 3c; *p* < 2.2×10^‒16^; Fisher’s exact test).

**Fig. 3.**
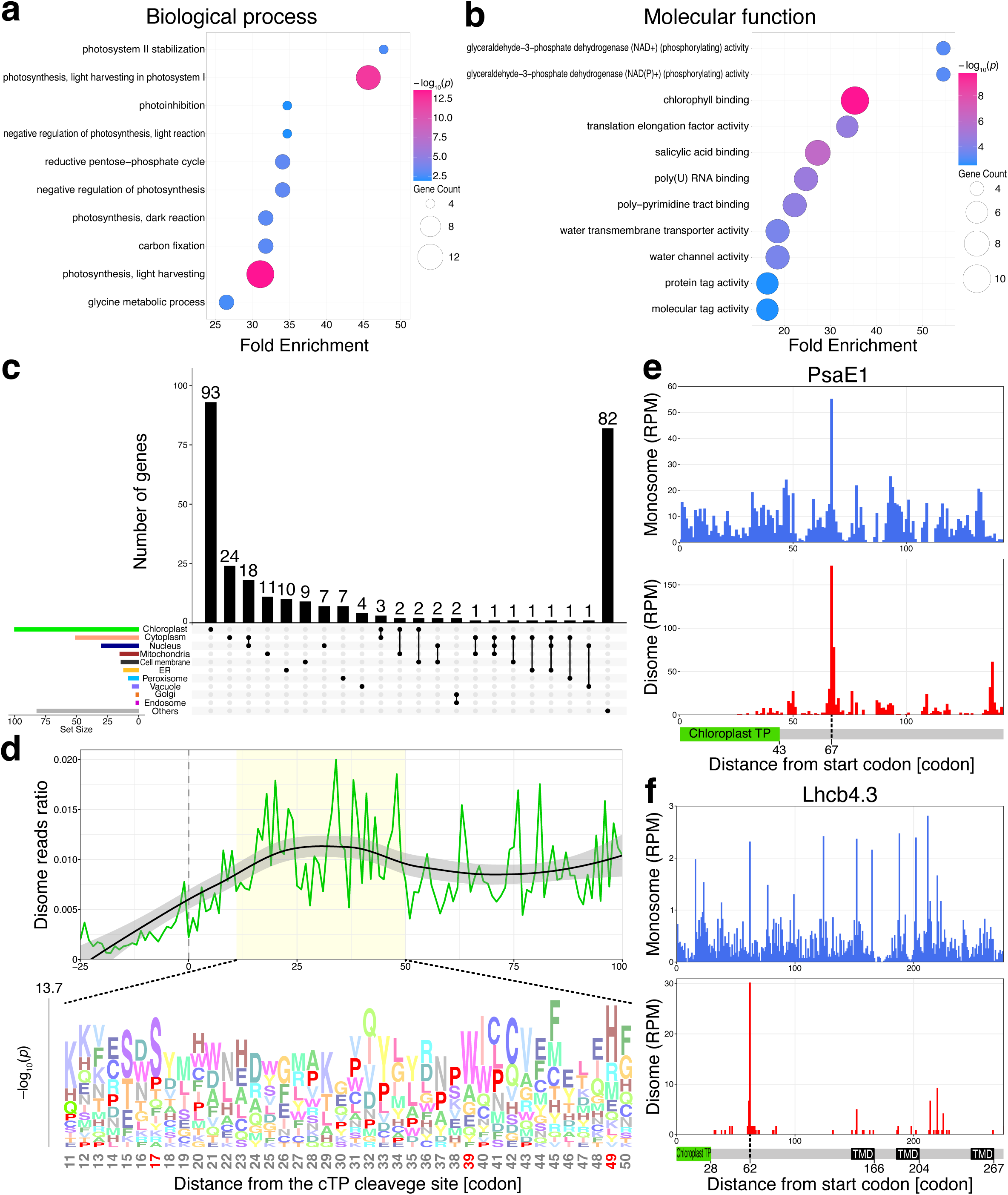
Ribosome stalling coincides with the emergence of a part or all of the transit peptide from the ribosome exit tunnel. **a**,**b**, Gene Ontology (GO) analysis of the 283 genes exhibiting CDS stalling. Bubble plots show enriched biological processes (**a**) and molecular functions (**b**). Bubble size represents the number of genes associated with each GO term, and bubble color represents –log_10_(*p*-value). Fold enrichment is plotted on the x-axis. **c**, Subcellular localization of proteins encoded by genes exhibiting ribosome stalling within CDSs. The UpSet plot visualizes the pattern of organelle localization based on UniProt annotations. Horizontal bars (left) indicate the total number of proteins annotated for each organelle, and vertical bars (top) indicate the total number of proteins in each intersection. “Others” includes proteins with unclassified or unknown localization. **d**, Metagene analysis of ribosome stalling sites relative to the predicted chloroplast transit peptide (cTP) cleavage site. The upper panel (green line) shows the disome footprint distribution within the CDS of chloroplast-localized genes satisfying the stalling criteria. Position 0 corresponds to the cTP cleavage site predicted by TargetP-2.0^37^. The black line represents the fitted curve, and the gray shading indicates the 95% confidence interval. The region 11–50 residues downstream of the cleavage site is highlighted in yellow. The lower panel shows the amino acid motif analysis within the yellow-highlighted region. Enrichment scores were calculated against all 40-amino-acid windows of nuclear-encoded proteins. Sequence logos were generated using *k*pLogo^56^, with significantly enriched positions highlighted in red. **e**,**f**, Footprint distributions along chloroplast-localized transcripts: *PsaE1* (**e**) and *Lhcb4.3* (**f**). A-site positions for monosome (upper; blue bars) and the leading ribosome in a disome (lower; red bars) are shown. Schematics below each plot depict the full-length protein (gray line), including the cTP (green) and TMDs (black). RPM: reads per million mapped reads.

Because nuclear-encoded chloroplast proteins are synthesized with N-terminal transit peptides that mediate organelle targeting, we examined whether stalling sites exhibit positional bias relative to these sequences. Transit peptide cleavage sites were predicted using TargetP-2.0^37^, and disome footprints from the 100 chloroplast-localized genes were aligned relative to these predicted sites, spanning 25 amino acids upstream to 100 amino acids downstream. This metagene analysis revealed a pronounced accumulation of disome footprints 20 to 40 amino acids downstream of the cleavage site (Fig. 3d, upper panel). Although this distance is comparable to the ∼30-amino-acids accumulation peaks observed downstream of start codons (Fig. 1e), the disome enrichment occurs well beyond the predicted transit peptide length (8–113 amino acids; median 52 amino acids). This observation argues against technical explanation related the minimal distance required for disome footprint detection and instead suggests that ribosomes preferentially stall after the nearly full-length transit peptide emerges from the ribosome exit tunnel. Within this downstream region, proline residues were particularly enriched in the 30–40 amino acid interval (Fig. 3d, lower panel), corresponding to the position at which the transit peptide is expected to have fully emerged from the ribosome exit tunnel. Consistent with this global trend, ribosome stalling in representative photosynthesis-related genes, including *PsaE1* and *Lhcb4.3*, occurred ∼30 amino acids downstream of the transit peptide C-terminus (Fig. 3e, f). Together, these findings indicate that ribosome stalling preferentially occurs immediately after transit peptide emergence from the ribosome exit tunnel, suggesting that translational pausing may promote efficient chloroplast localization.

To test this hypothesis, we generated a fusion construct containing the N-terminal region of *PsaE1* (from the start codon to the ribosome stalling site) fused to EGFP and a 3×FLAG tag (Fig. 4a; FL). We also generated three derivative mutants lacking the stalling region (Δstall), the transit peptide (ΔTP), or both (Δstall&TP). These constructs were transiently expressed in *Nicotiana benthamiana* leaves by agroinfiltration, and their subcellular localization was observed by assessing GFP fluorescence. In leaves expressing the FL construct, GFP fluorescence predominantly colocalized with chloroplast autofluorescence (Fig. 4b). In contrast, the Δstall construct showed markedly reduced overlap with chloroplast autofluorescence, resembling the localization pattern of the ΔTP and Δstall&TP constructs. Quantification of colocalization based on the relative GFP intensity—defined as the fraction of GFP signal overlapping with chloroplasts normalized to the mean GFP intensity across the field of view—revealed a significant reduction in chloroplast localization of the Δstall construct compared with the FL construct (Fig. 4c). A similar reduction was observed for *Lhcb4.3* constructs (Extended Data Fig. 4a, b), indicating that this effect is not gene-specific.

**Fig. 4.**
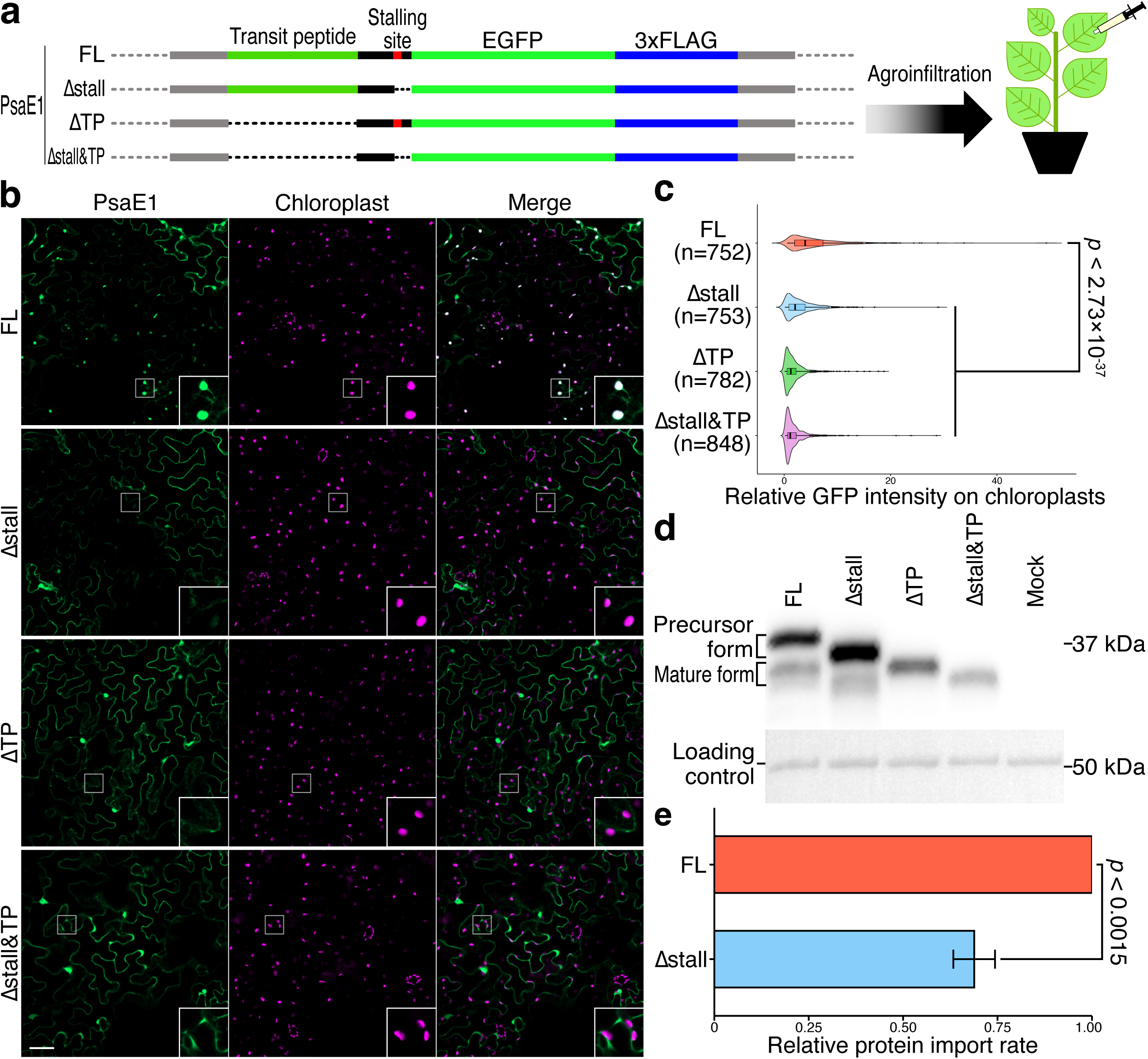
Ribosome stalling facilitates efficient chloroplast localization of proteins. **a**, Schematic workflow of transient expression in *N. benthamiana* leaves using Agrobacterium infiltration. Four PsaE1 constructs were tested: FL, Δstall, ΔTP, and Δstall&TP. Each construct contains EGFP and a C-terminal 3×FLAG tag. **b**, Confocal images of *N. benthamiana* leaves transiently expressing each construct. Observations were made 24 hours post-infiltration. Signals from PsaE1-EGFP (green), chloroplast autofluorescence (magenta), and merged channels are shown. Insets show magnified views of boxed regions. Scale bar, 50 µm. **c**, Quantification of chloroplast colocalization efficiency. Violin and box plots show the relative GFP intensity on chloroplasts, normalized to the mean GFP intensity of the entire field of view. n represents the number of chloroplasts analyzed. Statistical significance was determined using the Kruskal–Wallis test followed by Dunn’s multiple comparison test. **d**, Western blot analysis of leaf lysates harvested 24 h post-infiltration. PsaE1-EGFP fusion proteins were detected with anti-FLAG antibody. Ponceau-S staining is shown as a loading control. **e**, Quantification of relative protein import rate (mature/precursor ratio), normalized to the FL construct. Data represent mean ± s.d. Statistical comparison was performed using paired t-test.

Nuclear-encoded chloroplast proteins are synthesized as precursor forms bearing an N-terminal transit peptide that is cleaved upon import^38,39^. To directly assess chloroplast import, we monitored transit peptide processing by immunoblotting. In *N. benthamiana* leaves transiently expressing the constructs, both FL and Δstall variants produced two distinct bands corresponding to the precursor and mature forms, with the mature form comigrating with the ΔTP (Fig. 4d). Densitometric quantification of the immunoblot bands revealed a significant reduction in the mature-to-precursor ratio in the Δstall construct compared with FL (Fig. 4e), despite comparable overall protein expression levels. These results demonstrate that ribosome stalling enhances efficient chloroplast import at the stage when the transit peptide has emerged from the ribosome exit tunnel.

In this study, we show that translation of mRNAs encoding chloroplast-localized proteins is frequently accompanied by ribosome stalling and that this stalling enhances the efficiency of protein targeting to chloroplasts (Extended Data Fig. 5). Nuclear-encoded chloroplast-localized proteins have traditionally been thought to be imported predominantly via post-translational translocation, in which translation is completed in the cytoplasm before N-terminal transit peptides are recognized by cytosolic chaperones and delivered to the TOC–TIC complexes^40–42^. However, how translational dynamics are coordinated with chloroplast targeting has remained unclear^40,43,44^.

Ribosome stalling and local translational slowdown are known to promote nascent chain folding and facilitate chaperone recruitment as polypeptides emerge from the ribosome exit tunnel^45–47^. In line with this principle, our findings suggest that ribosome stalling on transcripts encoding chloroplast-associated proteins creates a temporal window that enhances transit peptide accessibility and promotes targeting competence.

Evidence from animal systems further indicates that ribosome stalling can contribute to organelle targeting. In mitochondria, which share an endosymbiotic origin with chloroplasts, a subset of proteins undergoes co-translational translocation^48–53^. In light of these observations, our data support a model in which chloroplast-localized proteins may similarly utilize a co-translational translocation pathway that operates alongside the canonical post-translational pathway (Extended Data Fig. 5). Supporting this possibility, previous studies in *Chlamydomonas* have shown that mRNAs encoding photosynthetic proteins accumulate at the chloroplast periphery and that cytoplasmic ribosomes associate with chloroplasts during active translation^54,55^.

Collectively, these results position ribosome stalling as a regulatory mechanism that temporally coordinates transit peptide exposure with chloroplast targeting, potentially coupling translation dynamics to organelle localization.

**Extended Data Fig. 1.**
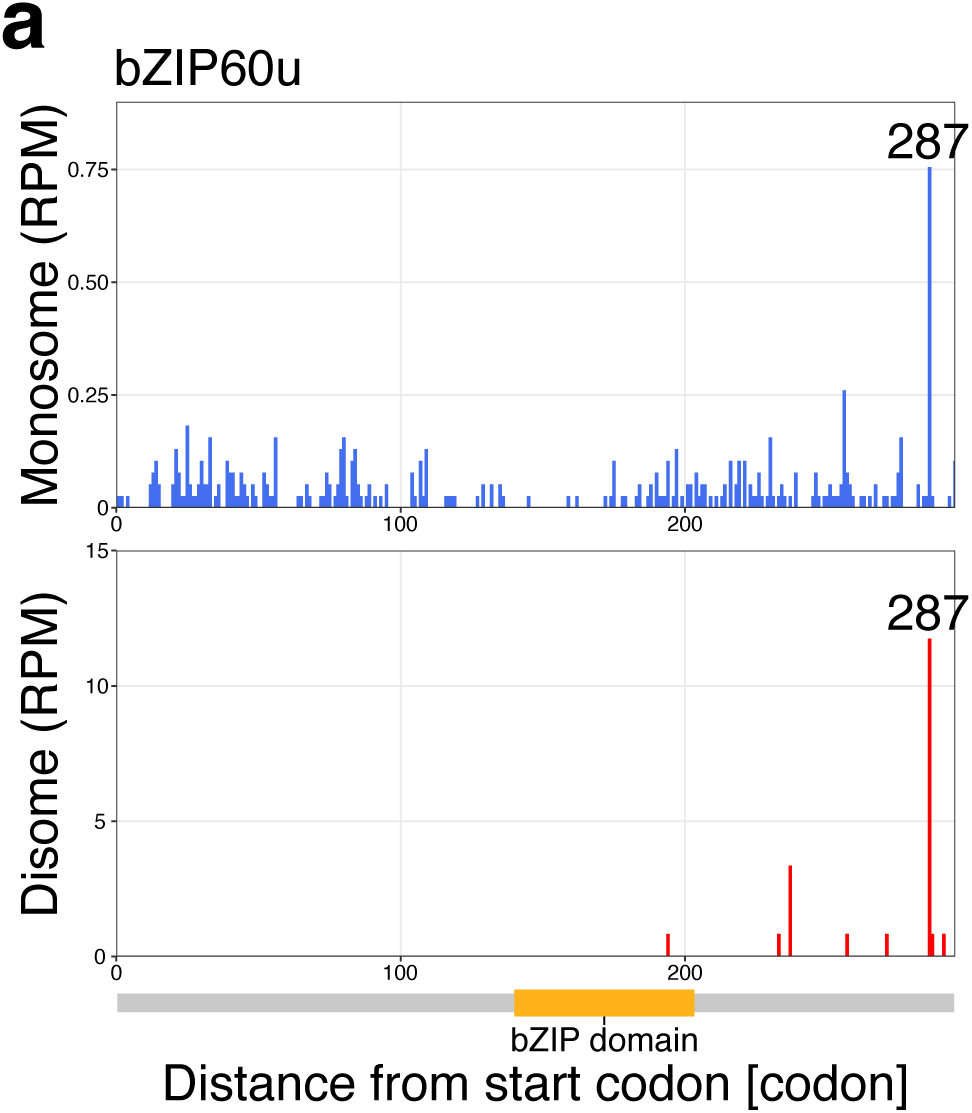
Benchmarking analysis using established ribosome stalling genes. Footprint distributions along nuclear-encoded bZIP60u transcript. A-site positions are plotted for monosome (upper; blue bars) and the leading ribosomes in a disome (lower; red bars). The established stalling site at Lys287 (287) is indicated in both panels. A schematic of the full-length protein is shown below (gray line), with the bZIP domain highlighted in orange. RPM: reads per million mapped reads.

**Extended Data Fig. 2.**
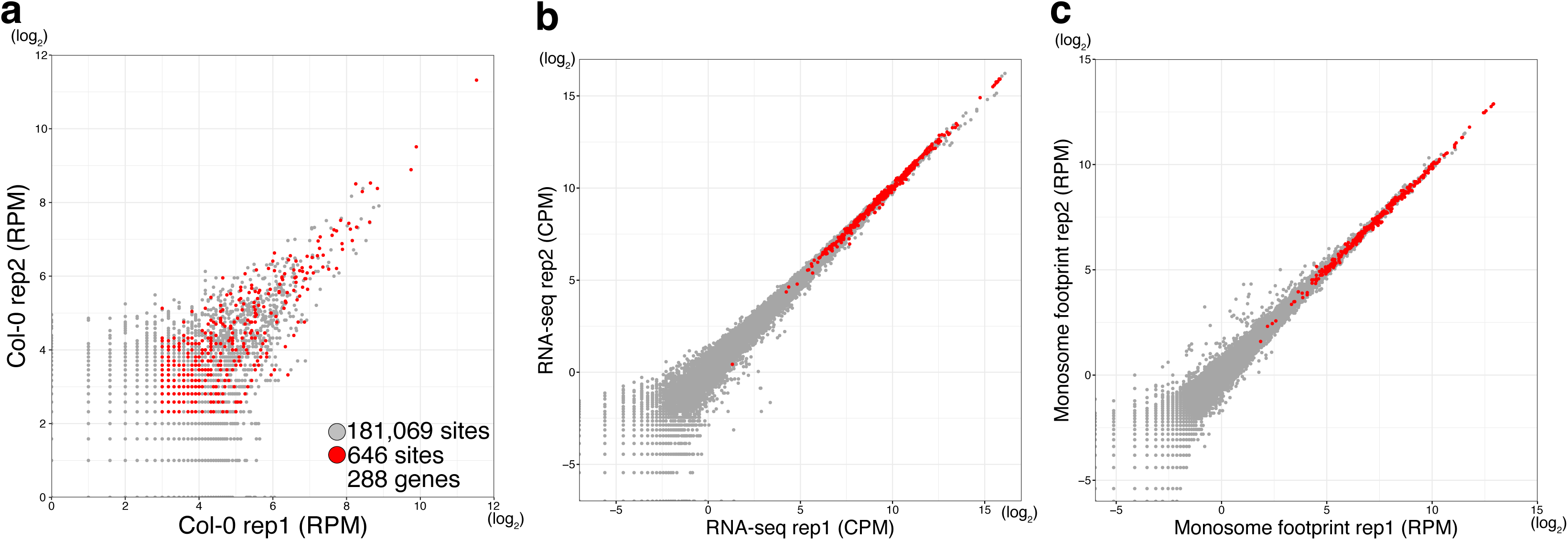
Transcriptional and translational levels of genes exhibiting ribosome stalling. **a**, Scatter plot of disome footprints by each replicate mapped to all nuclear-encoded transcripts annotated in TAIR10. Each point represents the first nucleotide of the A-site of the leading ribosome in a disome. Gray dots indicate all detected positions, and red dots indicate positions that satisfy the stalling criteria. **b**,**c**, Scatter plots comparing biological replicates of RNA-seq (**b**) and monosome footprints (**c**) data for all nuclear-encoded transcripts annotated in TAIR10. Each point represents an individual transcript; gray dots indicate all detected transcripts, and red dots indicate the transcripts corresponding to genes with prominent stalling identified in the transcriptome-wide analysis. Each axis shows log_2_-transformed values. CPM: counts per million reads. RPM: reads per million mapped reads.

**Extended Data Fig. 3.**
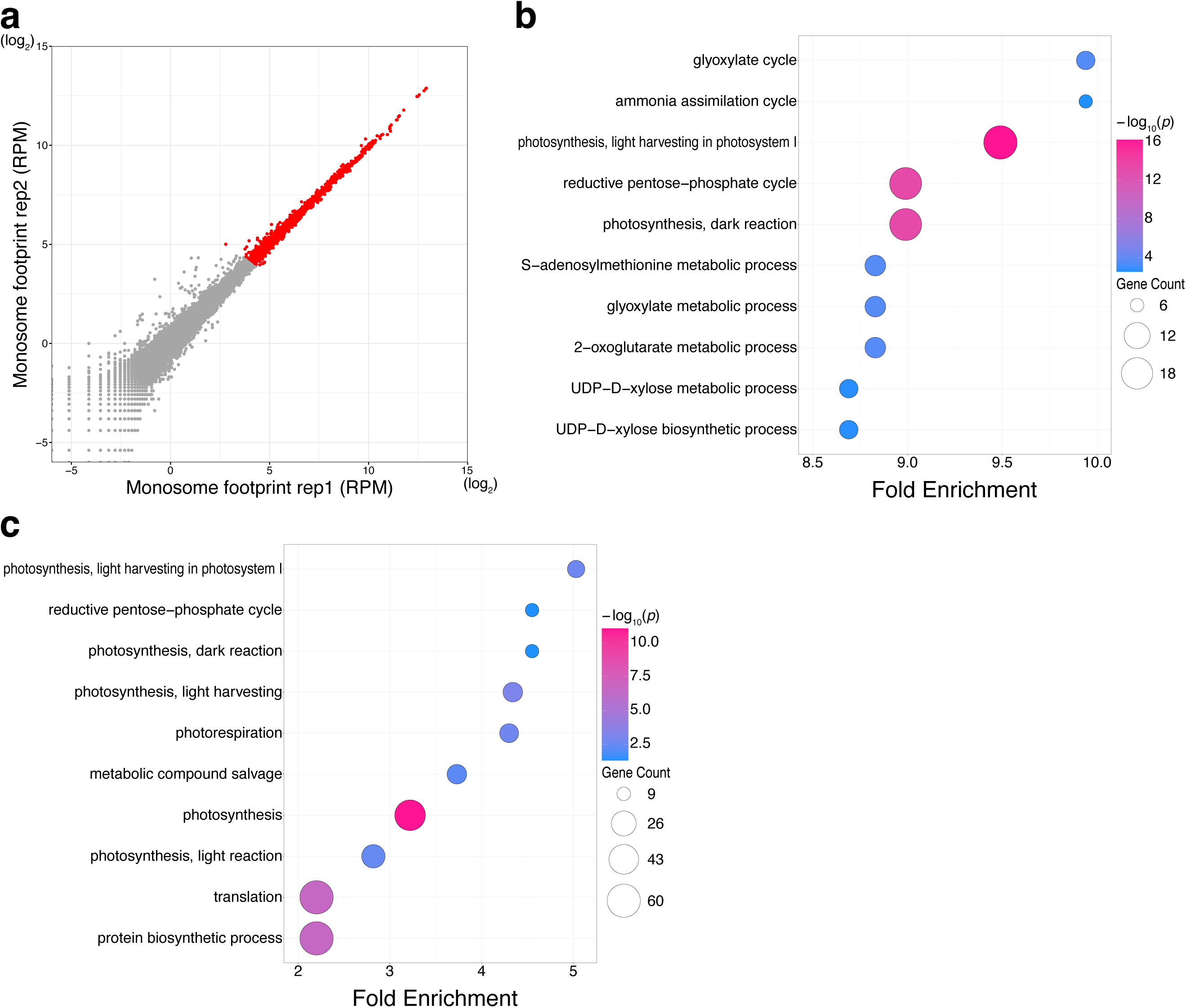
GO analyses of transcripts with highly accumulated monosome footprints. **a**, Scatter plots comparing biological replicates of monosome footprint data for all nuclear-encoded transcripts annotated in TAIR10. Each point represents an individual transcript; gray dots indicate all detected transcripts, and red dots indicate the top 10% monosome footprints-accumulated transcripts. Each axis shows log2-transformed values. RPM: reads per million mapped reads. **b**,**c**, GO analysis of the top 10% of transcripts with the highest monosome footprint accumulation (**b**) and the 283 genes exhibiting CDS stalling compared with this highly translated background (**c**). Bubble plots show enriched biological processes. Bubble size represents the number of genes associated with each GO term, and bubble color represents – log_10_(*p*-value). Fold enrichment is plotted on the x-axis.

**Extended Data Fig. 4.**
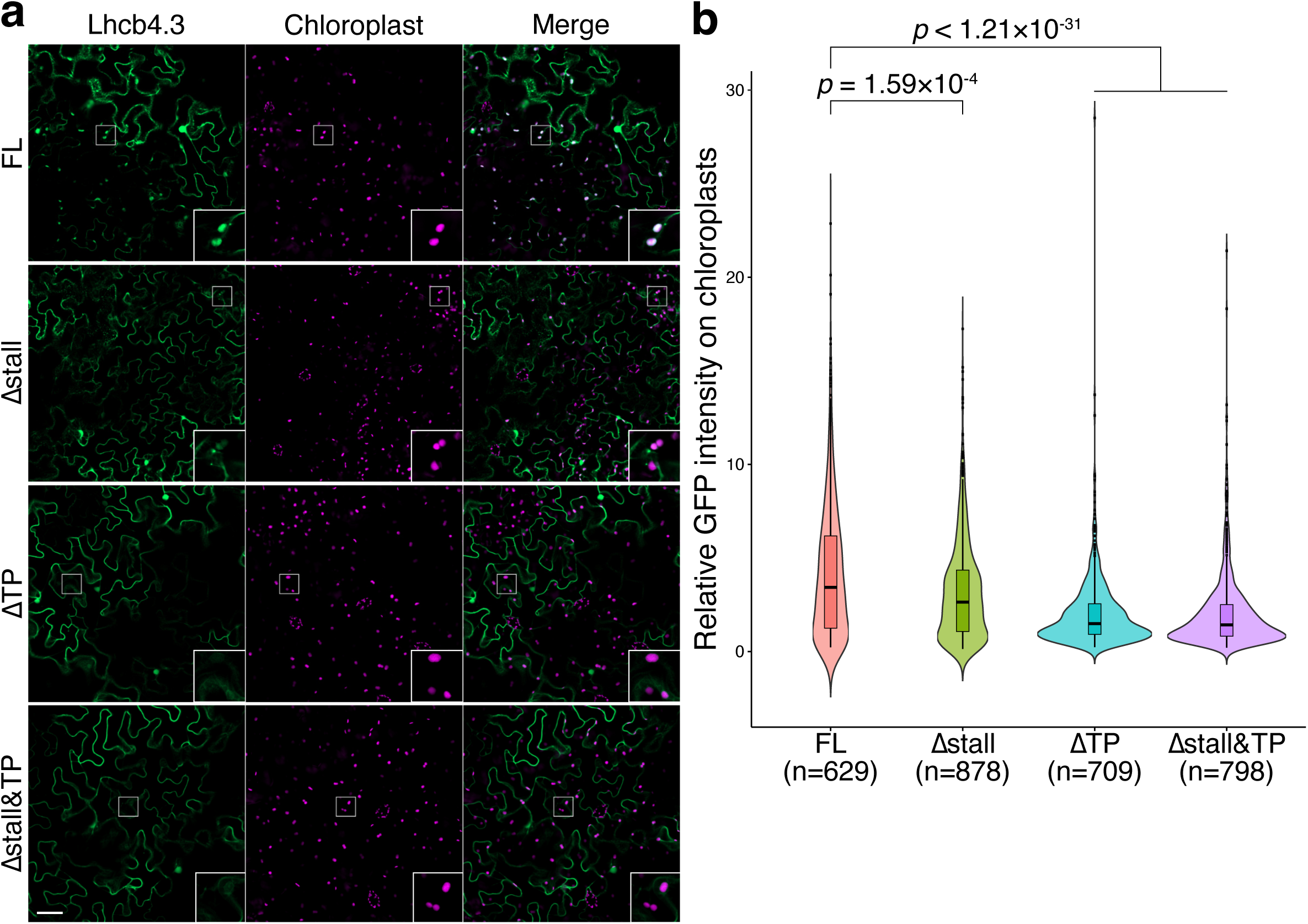
Localization analysis of Lhcb4.3 in *N. benthamiana* leaves. **a**, Confocal images of *N. benthamiana* leaves transiently expressing each Lhcb4.3 construct. Observations were made 24 h post-infiltration. Signals from Lhcb4.3-EGFP (green), chloroplast autofluorescence (magenta), and merged channels are shown. Insets show magnified views of boxed regions. Scale bar, 50 µm. **b**, Quantification of chloroplast colocalization efficiency. Violin and box plots show the relative GFP intensity on chloroplasts, normalized to the mean GFP intensity of the entire field of view. n represents the number of chloroplasts analyzed. Statistical significance was determined using the Kruskal–Wallis test followed by Dunn’s multiple comparison test.

**Extended Data Fig. 5.**
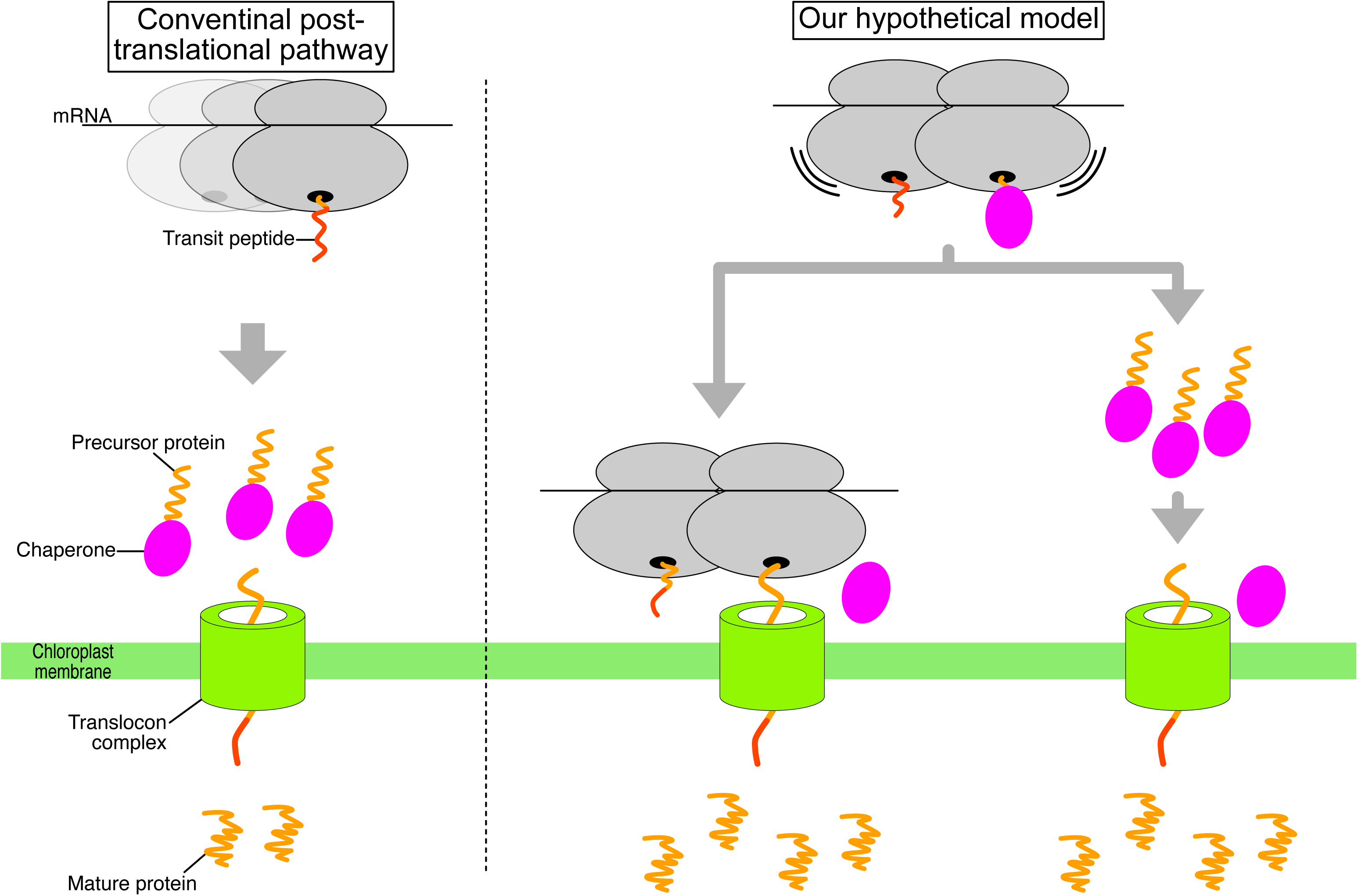
Model for the localization of chloroplast-localized proteins. Left: The conventional chloroplast targeting model, in which translation is completed in the cytosol and the precursor protein is subsequently delivered to the chloroplast surface for import through the translocon complex. Right: The ribosome stalling-mediated chloroplast targeting model, in which ribosome stalling facilitates efficient targeting and import of nascent proteins into the chloroplasts.

## Methods

### Plant materials and growth conditions

*A. thaliana* Columbia-0 (Col-0) was used as the wild type. Seeds were surface-sterilized with 70% EtOH at room temperature for 2 min, followed by a 10% sodium hypochlorite solution for 5 min, and washed five times with sterile water. Sterilized seeds were sown on filter paper (Whatman No.2) placed on Murashige and Skoog (MS) agar plates (1×MS salt, 1% sucrose, 1% agar, pH 5.7). After stratification in the dark at 4 °C for 3 days, the plates were incubated at 22°C for 3 days under continuous LED light (LC-LED450W, TAITEC).

*N. benthamiana* seeds were sown in soil pots and watered with Hyponex (Hyponex Japan) diluted 1:2,000. Plants were grown under long-day conditions (16-h light/ 8-h dark) at approximately 22 °C for 6 weeks before infiltration.

### Plasmid construction

The pMDC32 vector used in this study has been described previously^57^. The cDNA fragments of *PsaE1* (AT4G28750) and *Lhcb4.3* (AT2G40100) were amplified using cloning primers (Supplementary Table 1). DNA fragments corresponding to the FL and deletion mutants were amplified using specific primers (Supplementary Table 1) and assembled with the PCR-amplified pMDC32 vector fragments using the NEBuilder HiFi DNA Assembly (E2621, New England Biolabs).

### Library preparation for disome profiling

Disome profiling libraries were prepared following previously described protocols^17,24^, with some modifications for *A. thaliana* seedlings. Following RNA digestion with RNase I (N6901K, LGC, Biosearch Technologies), RNA fragments were pelleted by centrifugation through sucrose cushion buffer (100 mM Tris-HCl (pH 7.5), 40 mM KCl, 20 mM MgCl_2_, 1 mM DTT, 300 µg/mL Cycloheximide, 1 M Sucrose, and 20 U/mL SUPERase•In (Thermo Fisher Scientific)). The pellet was resuspended in Pellet lysis buffer (100 mM Tris-HCl (pH 7.5), 80 mM KCl, 5 mM EDTA, 1 mM DTT, 1% Triton X-100, and 6 U/mL SUPERase•In). The suspension was transferred to an Amicon Ultra 0.5 mL centrifugal filter device (Millipore) and centrifuged at 14,000 × *g* for 10 min at 4 °C^58^. After removing the filter, TRIzol LS reagent (Thermo Fisher Scientific) was added to the flow-through, and RNA fragments were purified using the Direct-zol RNA MicroPrep Kit (Zymo Research) according to the instructions. Purified RNA fragments ranging from 40 to 80 nt were gel-extracted for disome profiling library preparation.

### Disome profiling data analysis

Disome profiling data were processed as previously described^22,23^. Adaptor sequences were trimmed using Fastp^59^. After removing adaptor sequences and reads mapping to non-coding RNA (including rRNA and tRNA sequences), the remaining reads were aligned to the TAIR10 *A. thaliana* reference genome using STAR (version 2.7.0a)^60^. Disome footprints with lengths of 54–58 nt were selected for analysis. The A-site position for each disome footprint was defined by applying an offset from the 5′ end aligning to the start codons of the transcripts. The A-site offsets are 41 for 54 nt, 42 for 55 nt, 43 for 56 nt, 44 for 57 nt, and 45 for 58 nt.

### Monosome profiling and RNA-seq data analysis

Monosome profiling data were reanalyzed using previously published datasets^17^ (DRA010034), employing the reported footprint size and A-site offsets.

RNA-seq data were also processed as previously described^17^.

### Bioinformatics analyses

GO analysis was performed using PANTHER 19.0^61,62^.

The amino acid motifs associated with ribosome stalling sites were visualized using *k*pLogo^56^ (http://kplogo.wi.mit.edu/). To classify stalling motif patterns, ribosome stalling sites were clustered using GibbsCluster-2.0^63^ and subsequently visualized with *k*pLogo.

### Transient expression in *N. benthamiana* leaves

The FL and deletion mutant constructs were transformed into *Agrobacterium tumefaciens* strain GV3101^64^. The *Agrobacterium* suspensions (adjusted optical density at 600 nm (OD600) = 0.2 (each *Lhcb4.3* construct) or 0.5 (each *PsaE1* construct)) were infiltrated into *N. benthamiana* leaves using a needleless 1 mL syringe (Terumo). The leaves were harvested at 24 h post-infiltration.

### Confocal microscopy and image analysis

Infiltrated leaves were observed using an LSM980 confocal laser scanning microscope (Carl Zeiss) equipped with a 20× dry objective lens. Confocal images were acquired for each construct at a resolution of 1,024×1,024 pixels using bidirectional scanning with a speed of 1. Image analysis was performed using Fiji software^65^. Chloroplast areas were automatically defined using the threshold tool, and GFP intensities overlapping with these areas were measured. For the representative images shown in Fig. 4b and Extended Data Fig. 3a, brightness and contrast were adjusted for improved visualization.

### Protein extraction and Western blotting

Total protein was extracted from infiltrated leaves in 100 μL of Homogenization buffer (0.33 M sorbitol, 1 mM MgCl_2_, 20 mM MES, 0.1% (w/v) Ficoll, and 0.5 mM Tricine) using a homogenizer (BioMasher II, nippi). Samples were mixed with 6× SDS-PAGE sample buffer (280 mM Tris-HCl (pH 7.5), 30% (v/v) glycerol, 10% (v/v) SDS, 9.3% (w/v) DTT, and 0.02% (w/v) bromophenol blue) to a final concentration of 1.2× and boiled at 95 °C for 5 min. Boiled samples were separated by SDS-PAGE and then transferred onto a PVDF membrane (IPVH00010, Millipore). The membrane was blocked with 1% (w/v) nonfat dried milk in TBST for 10 min. The membrane was incubated with anti-DDDDK-tag mAb (diluted at 1:8,000; M185-3L, MBL) as the primary antibody, followed by HRP-conjugated anti-mouse IgG (H+L) pAb (diluted at 1:8,000; 330, MBL) as the secondary antibody. Signals were detected using a chemiluminescent substrate (34580, Thermo Fisher Scientific) and imaged using a FUSION FX (Vilber).

## Statistical analysis

All statistical analyses were performed using R (version 4.2.2). The statistical tests performed on experimental data and the sample sizes are noted in the figure or legends. All data points are derived from biological replicates.

## Data availability

The disome profiling data generated in this study (DRA026166) were deposited in the National Center for Biotechnology Information (NCBI) database. This study also used previously deposited ribosome profiling data^17^ (DRA010034).

## Acknowledgements

We thank Yuichi Shichino and Mari Mito for technical assistance with ribosome profiling; Keisuke Shoji for kind advice with NGS data analyses. We also thank all members of the Tomari laboratory and Iwakawa laboratory for discussions and critical comments on the manuscript. This work was supported by JSPS KAKENHI (grant numbers 24KJ0577 to R.K., 23H02412 to H.-o.I.), JST FOREST (grant JPMJFR204O to H.-o.I.), JST PRESTO (grant JPMJPR18K2 to H.-o.I.), and RIKEN (Pioneering project to S.I.). This study uses the HOKUSAI SailingShip Supercomputer facility at RIKEN for computation.

## Author Contributions

R.K. and H.-o.I. designed the project. R.K. performed ribosome profiling and associated bioinformatics analyses under the supervision of S.I. R.K. performed biochemical experiments and additional bioinformatics analyses under the supervision of Y.T. and H.-o.I. R.K. and H.-o.I. wrote the manuscript with input from all authors. All authors discussed the results and approved the final version of the manuscript.

## Notes

### Competing Interest Statement

The authors have declared no competing interest.

