## Supplementary Table 1 for "Ribosome stalling facilitates chloroplast targeting of nuclear-encoded proteins"

| Name | Sequence | Purpose |
|---|---|---|
| pBYL_PsaE1_mRNA | TTGCATGCCTGCAGGTCGACTCTAGAGTAATGTACACGTGGCAGATCTCAAGTGGAGGGAAACGATTCGTTTAACCCAAAACATCTC | forward primer for *PsaE1* full length cDNA fragment |
| PsaE1_mRNA_pBYL | ATACACCCAAAAGTCTCAAGCTGGCGCGCCAAGAGAGCTAACCATGAGTAGAAAGAGACTTTTAACTGAATTTTCCAAACACATTCTGTG | reverse primer for *PsaE1* full length cDNA fragment |
| pMDC_Lhcb4.3_5´UTR_Fw_1 | AGGAAGTTCATTTCATTTGGAGAGGAAAAGAAAGAAAATACCTTAAATGCAGACACCTACCATAAGGACGAAAG | forward primer for *Lhcb4.3* full length cDNA fragment |
| Lhcb4.3_3´UTR_pMDC_Rev_1 | CGGTGGGCGGCCGCTCTAGAACTAGTCACGGTATATTAGTTTAGAACATAAACAATAAATACTTCGATATTTAT | reverse primer for *Lhcb4.3* full length cDNA fragment |
| pMDC32_Fw_1 | GCATCCGCTTACAGACAAGC | forward primer for pMDC32 1st half fragment |
| pMDC32_Rev_1 | ACGAGTGTTTACAGCGATAATGCTAGCGAATCAACAGTGAAGAACTTGC | reverse primer for pMDC32 1st half fragment |
| pMDC32_Fw_2 | CTAGTTCTAGAGCGGCCGC | forward primer for pMDC32 2nd half fragment |
| pMDC32_Rev_2 | GAGACGGTCACAGCTTGTCTG | reverse primer for pMDC32 2nd half fragment |
| EGFP_Fw | ATGGTGAGCAAGGGCGAGG | forward primer for EGFP |
| 3×FLAG_pMDC32_Rev | GTGGCGGCCGCTCTAGAACTAGTCATCACTTGTCATCGTCATCCTTG | reverse primer for 3×FLAG tag with pMDC32 overlap |
| 5´UTR_PsaE1_Fw | CTAGCATTATCGCTGTAAACACTCGTATGGCGATGACGACAGC | forward primer for FL-*PsaE1* with pMDC32 overlap |
| PsaE1_stall_site_EGFP_Rev | ACAGCTCCTCGCCCTTGCTCACCATTTTGGTGGCAGTAGCTCCA | reverse primer for FL-*PsaE1* with EGFP overlap |
| 5´UTR_PsaE1_ΔTP_Fw | GCATTATCGCTGTAAACACTCGTATGGCAGCCGAAGATCCTGC | forward primer for ΔTP-*PsaE1* with pMDC32 overlap |
| PsaE1_Δstall_site_EGFP_Rev | ACAGCTCCTCGCCCTTGCTCACCATAGCAGCGGCAGCTGCCGG | reverse primer for Δstall-*PsaE1* with EGFP overlap |
| 5´UTR_Lhcb4.3_Fw | TAGCATTATCGCTGTAAACACTCGTATGGCTACCACCACTGCAGC | forward primer for FL-*Lhcb4.3* with pMDC32 overlap |

| | | |
|---|---|---|
| Lhcb4.3_stall_site_EGFP_Rev | ACAGCTCCTCGCCCTTGCTCACCATCCATTCCGGCGGGTTTGC | reverse primer for FL-*Lhcb4.3* with EGFP overlap |
| 5´UTR_Lhcb4.3_△TP_Fw | GCATTATCGCTGTAAACACTCGTATGCGGTTCGGGTTCAGTTTCG | forward primer for ΔTP-*Lhcb4.3* with pMDC32 overlap |
| Lhcb4.3_△stall_site_EGFP_Rev | GCTCCTCGCCCTTGCTCACCATGAACCAAACTAGTCTGTCTCCATCG | reverse primer for Δstall-*Lhcb4.3* with EGFP overlap |
